# Oligomerization state of the functional bacterial twin arginine translocation (Tat) receptor complex

**DOI:** 10.1101/2021.11.01.466846

**Authors:** Ankith Sharma, Rajdeep Chowdhury, Siegfried M. Musser

## Abstract

The twin-arginine translocation (Tat) system transports folded proteins across bacterial and plastid energy transducing membranes. Ion leaks are generally considered to be mitigated by the creation and destruction of the translocation conduit in a cargo-dependent manner, a mechanism that enables tight sealing around a wide range of cargo shapes and sizes. In contrast to the variable stoichiometry of the active translocon, the oligomerization state of the receptor complex is considered more consistently stable, but has proved stubbornly difficult to establish. Here, using a single molecule photobleaching analysis of individual inverted membrane vesicles, we demonstrate that Tat receptor complexes are tetrameric in native membranes with respect to both TatB and TatC. This establishes a maximal diameter for a resting state closed pore. A large percentage of Tat-deficient vesicles explains the typical low transport efficiencies observed. This individual reaction chamber approach will facilitate examination of the effects of stochastically distributed molecules.

## INTRODUCTION

The Tat machinery has the unusual capacity to transport folded proteins across energetic membranes without collapsing ion gradients^1–6^. It accomplishes this without requiring any nucleoside triphosphates (NTPs) and with the unique requirement that a proton motive force (pmf) is essential for transporting all substrates^3,7,8^. Despite the seemingly complex physical problem of conveying a large object across a membrane barrier while maintaining the permeability barrier, a minimal Tat system requires only two membrane proteins, TatA and TatC, e.g., as found in some gram-positive bacteria and archaea^9,10^. Nonetheless, the best-studied Tat systems are found in *Escherichia coli* and in plant thylakoids, both of which additionally contain TatB, a TatA-like membrane protein. Though *E. coli* also contains a fourth protein, TatE, which is homologous to TatA and TatB^6,11,12^, *E. coli* TatABC comprise a common minimal system^6,7,13–18^. An early step in the transport cycle is recruitment of substrates with a twin-arginine signal peptide (RRXFLX consensus motif) by the Tat receptor complex^6,19^. However, transport cannot occur without a TatA-rich assembly, typically described as resulting from recruitment of TatA to the receptor-substrate complex^20–22^. This active translocon is proposed to have a variable composition, allowing it to accommodate a range of mature domain sizes and shapes^23–25^. In contrast, the receptor complex is generally assumed to have a fixed composition, yet it has been surprisingly difficult to establish its oligomerization state^22,26–31^.

TatC has six transmembrane helices that generate a glove-like groove within the membrane and forms the bulk of the signal peptide binding pocket^32,33^. Membranes containing TatB and TatC are sufficient for recruiting Tat substrates, establishing that these proteins form the core of the receptor complex^17,34,35^, and these are generally agreed to be present in a 1:1 ratio^22,36,37^. Whether TatA is a normal constituent of the receptor complex has been controversial; at present, the weight of evidence leans toward some TatA in the receptor complex^17,27–29,31^. More recently, it was discovered that TatA, TatB, and TatE occupy very similar positions on TatC^38^, suggesting that TatE is also part of the receptor complex, or that TatA takes the place of TatE when the latter is absent. Thus far, there is no evidence that any additional TatA and/or TatE within the receptor complex is in excess of one molecule per TatBC heterodimer. Thus, their presence would be insufficient to substantially increase the size of the receptor complex, although they could influence oligomerization state estimates. Due to the longstanding uncertainty on the presence of TatA and TatE in the receptor complex, its oligomerization state is typically discussed in terms of the number of TatBC heterodimers.

The receptor complex oligomerization state is fundamental to a mechanistic understanding of the transport mechanism. Considering that the receptor-bound signal peptide is inserted about halfway across the membrane^17^, a fundamental structural issue is whether the signal peptide binding site is exposed to the membrane lipid interior, which it must be for a monomeric receptor complex, or whether it could be protected within the interior of a higher-order receptor oligomer^39^. For lipid-exposed binding site models, recruitment of TatA could, in principle, generate a translocation conduit with minimal (if any) conformational changes within the receptor complex. In contrast, for interior binding site models, significant conformational changes appear necessary to rearrange to create a channel de novo, or to gate the channel open and closed, possibly also increasing the diameter of a resting, but blocked, pore^29,39–41^. Notably, higher-order oligomers could form larger ‘resting’ pores, although they present a more substantial challenge to seal against ion leaks.

The receptor complex oligomerization state gleaned from various studies includes a dimer, trimer, tetramer, heptamer and octamer, although trimer and tetramer models are favored more recently^22,26–31,36,37^. Large ~440-700 kDa complexes obtained upon detergent extraction^22,30,31,36,37,42^ can over-estimate oligomer size due to aggregation or co-purifying molecules, such as lipids. Crosslinking approaches^27–29^ can either under- or over-estimate complex size due to insufficient crosslinks or crosslinks to unrelated proteins. Co-evolution analysis requires inferring the correct structural model and multiple oligomeric forms may be equally viable^29^. While conflicting conclusions over oligomerization state can certainly point toward variable compositions as conditions change, we considered that direct measurement by molecular counting in the native membrane environment would provide a more robust result.

Here, we optimized the overproduction level such that at most a couple of Tat receptor complexes are recovered per inverted membrane vesicle (IMV). Assuming that the cell resuspension and IMV formation processes do not disrupt the Tat receptor oligomer, the receptor complex oligomerization state in its native lipid environment can be inferred from the distribution of the Tat proteins in IMVs. Using a molecular counting approach, we show that in IMVs from cells overproducing TatABC, both TatB and TatC are present as tetramers, the number of receptor complexes per IMV is ~1, and a large fraction of IMVs are devoid of Tat receptor complexes.

## RESULTS

### Experimental Design

An inherent underlying assumption when purifying membrane proteins with detergents is that the extraction process does not influence the composition of the protein complex. Verification that this is indeed so requires comparison with numerous controls. We avoided this complication by assaying the proteins present within native membrane vesicles. The formation of IMVs involves breaking the cell membrane into numerous pieces, which individually reseal to produce a membrane bilayer separating two aqueous compartments that in most cases is typologically inverted (the cytoplasmic leaflet is now on the outside)^7^. Thus, we expected that protein complexes present within the original cytoplasmic membrane would stochastically partition into the IMVs. Once the complexes were distributed in the IMVs, our counting assay does not rely on the protein assemblies remaining intact, merely that the individual polypeptides are expected to remain associated with the same IMV.

The overall goal of this study was to discern the number of TatB and TatC molecules present in the *E. coli* Tat receptor complex. We reasoned that by generating a low average number of receptor complexes per IMV via an appropriate expression level, the number of TatB or TatC molecules could be counted by single molecule photobleaching methods on individually assayed IMVs. Our primary analytical assumption was that histogram distributions summarizing the number of observed steps in photobleaching traces were expected to be dependent on the oligomer size. For example, a tetramer can be expected to give rise to spots with a majority of fluorescent intensities corresponding to four, eight, twelve, etc. molecules. Thus, we expected to be able to determine for the first time the average number and distribution of TatB and TatC molecules and oligomers per IMV. To achieve this, we separately C-terminally tagged TatB and TatC with the fluorescent protein mNeonGreen (mNeon)^43^ (**Supplementary Fig. 1**). IMVs were isolated from cells with Tat receptor complexes containing either TatB^mNeon^ or TatC^mNeon^, the membranes were labeled with the CellMask membrane stain, and the IMVs were then attached to microscope coverslips for photobleaching analysis (**Fig. 1**).

**Fig. 1.**
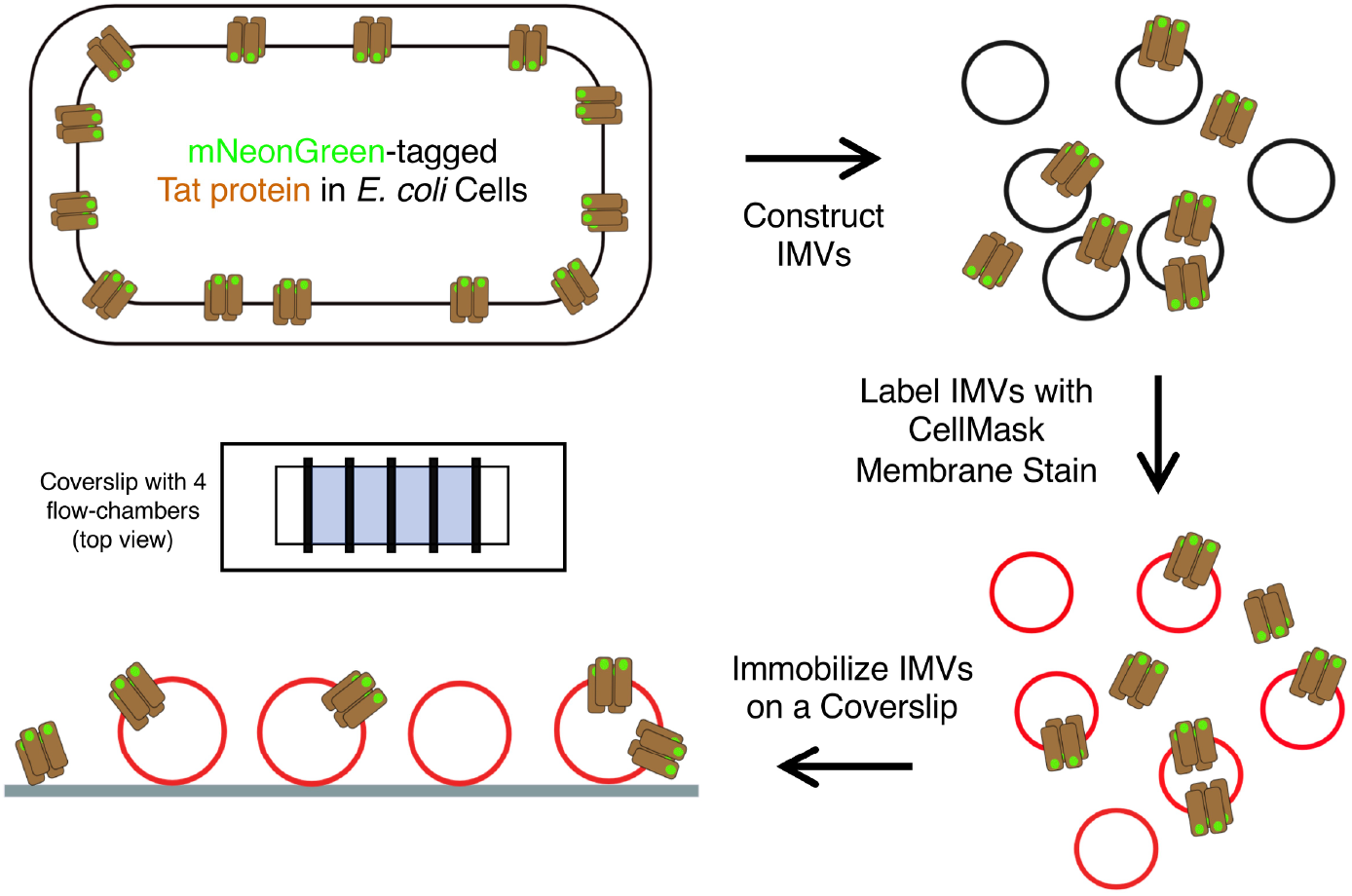
Experimental Strategy. IMVs were obtained from *E. coli* cells overproducing TatAB^mNeon^C or TatABC^mNeon^. After staining the membrane with CellMask, the IMVs were added to a flow-chamber constructed on a microscope coverslip. The adsorbed IMVs were washed before analysis.

### Suitability of IMVs with mNeon-labeled Tat Receptor Complexes for Molecular Counting Analysis

To ensure complete labeling of the TatB and TatC proteins, the TatABC proteins were overproduced from a plasmid in the deletion strain MC4100ΔTatABCDE (*a.k.a.* DADE)^44^. IMVs produced from such cells are considered Tat^++^ IMVs^45^. A 1 h induction time yielded the low average number of TatB^mNeon^ or TatC^mNeon^ molecules per IMV necessary for molecular counting. This is a significantly lower production level than for IMVs used for typical in vitro transport assays^7,18,46,47^. Not surprisingly, then, these IMVs containing TatB^mNeon^ or TatC^mNeon^ exhibited reduced transport efficiencies (**Fig. 2a**). No noticeable cleavage of the TatB^mNeon^ and TatC^mNeon^ proteins in IMV preparations was observed (**Supplementary Fig. 2**), indicating their suitability for molecular counting. The average diameters of TatB^mNeon^ and TatC^mNeon^ IMVs were 105 nm and 100 nm, respectively, as determined by dynamic light scattering (DLS; **Fig. 2c**). This small size indicates that individual IMVs should be visualized essentially as diffraction-limited spots when using light microscopy. Further, all fluorophores present in such IMVs are expected to be well-focused and simultaneously detectable in a single fluorescent spot. Individual complexes or molecular diffusion were not resolvable on the timescale of our experiments.

**Fig. 2.**
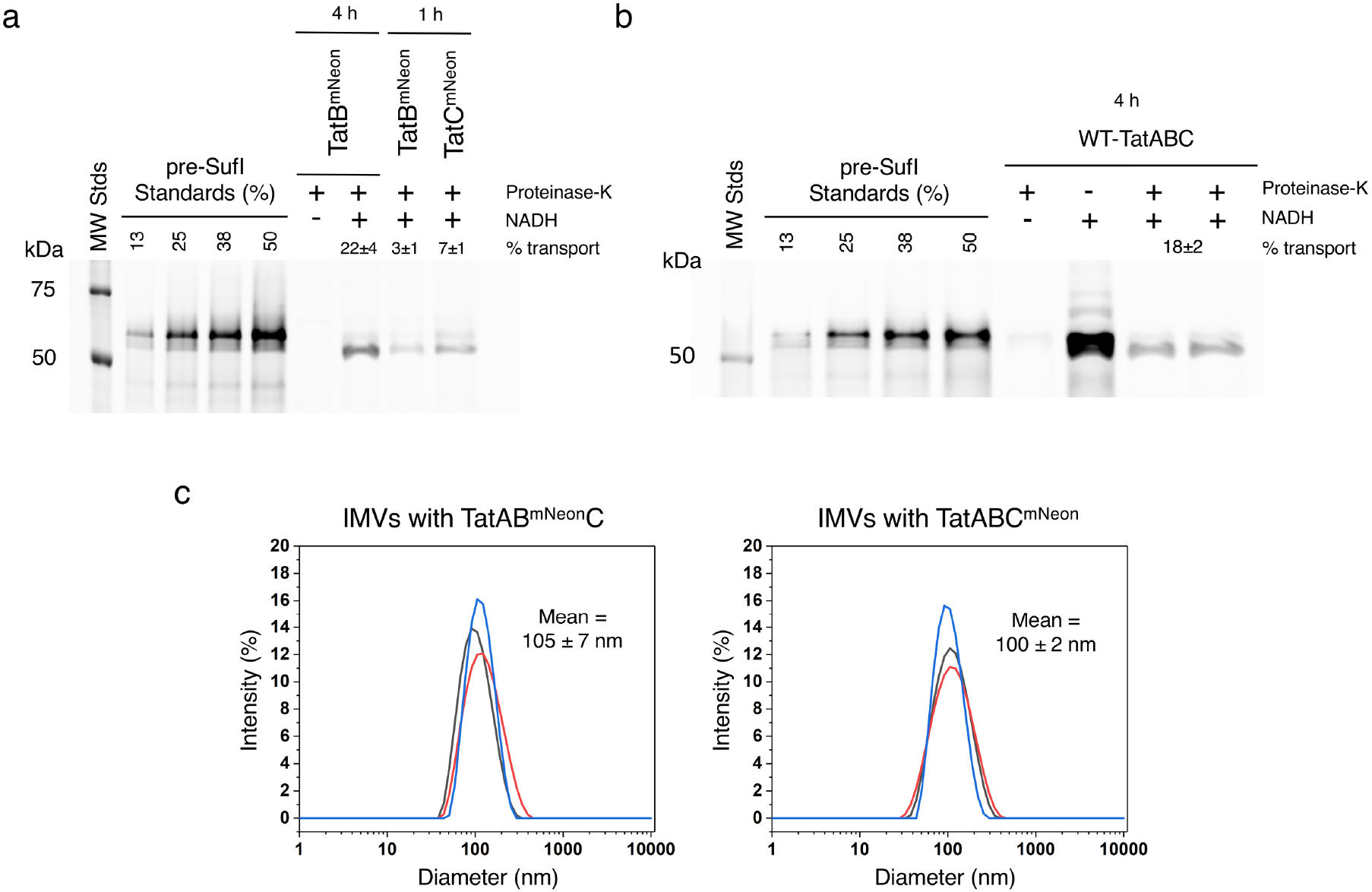
Transport Efficiency and Size of IMVs with TatAB^mNeon^C or TatABC^mNeon^. **a,b,** In vitro transport assays. The amount of SufI(IAC)^Alexa647^ (50 nM) transported into Tat^++^ IMVs was determined after 20 min at 37°C. Transport efficiency (% transport) was estimated from the amount of protease-protected mature-length protein compared with the amount of total added precursor protein (pre-SufI standards). The necessary pmf was generated by the addition of NADH (4 mM). A 4 h induction period yielded a similar transport efficiency for TatAB^mNeon^C (**a**) and wildtype TatABC (Tat^++^) IMVs (**b**). SDS-PAGE gels were analyzed by direct in-gel fluorescence imaging (λ_ex_ = 488 nm). A substantially lower transport efficiency was observed for a short (1 h) overproduction time (**a**), which was required to insure a low number of Tat receptor complexes per IMV. All transport efficiencies were calculated from duplicate assays. **c,** IMV size. Diameter distributions of three independent preparations of IMVs containing either TatAB^mNeon^C or TatABC^mNeon^, as determined by DLS. A consistent average diameter of ~100 nm was obtained.

### Photobleaching of IMVs Containing TatB^mNeon^ or TatC^mNeon^ IMVs

IMVs were adsorbed to plasma-treated coverslips at a dilution suitable for obtaining a sparse distribution of well-isolated fluorescence spots, which were visualized by narrow-field epifluorescence microscopy^48^. The mNeon and CellMask channels allowed independent identification of spots containing the labeled Tat protein and lipids, respectively (**Fig. 3a**). Colocalization analysis revealed that: (i) 18%-33% of IMVs contained an mNeon-tagged Tat protein and (ii) 30-65% of the mNeon-tagged Tat protein was associated with a lipid vesicle. While some larger, intensely fluorescent spots in the mNeon channel were observed (most of which were probably aggregates), the majority of spots were approximately diffraction-limited with variable intensities. Photobleaching traces of mNeon were collected for all of the smaller spots that co-localized with lipids, revealing a characteristic stepwise pattern with a variable number of steps (**Fig. 3b**). For comparison, and to establish an appropriate protocol for analysis and step counting, photobleaching traces for mNeon monomers, dimers and tetramers were obtained (**Supplementary Fig. 3** and **Fig. 4**). While fluorescence blinking was observed, the combination of intensity and observable steps yielded sufficient confidence in a manual approach to step counting. Consequently, all steps were counted by manual inspection.

**Fig. 3.**
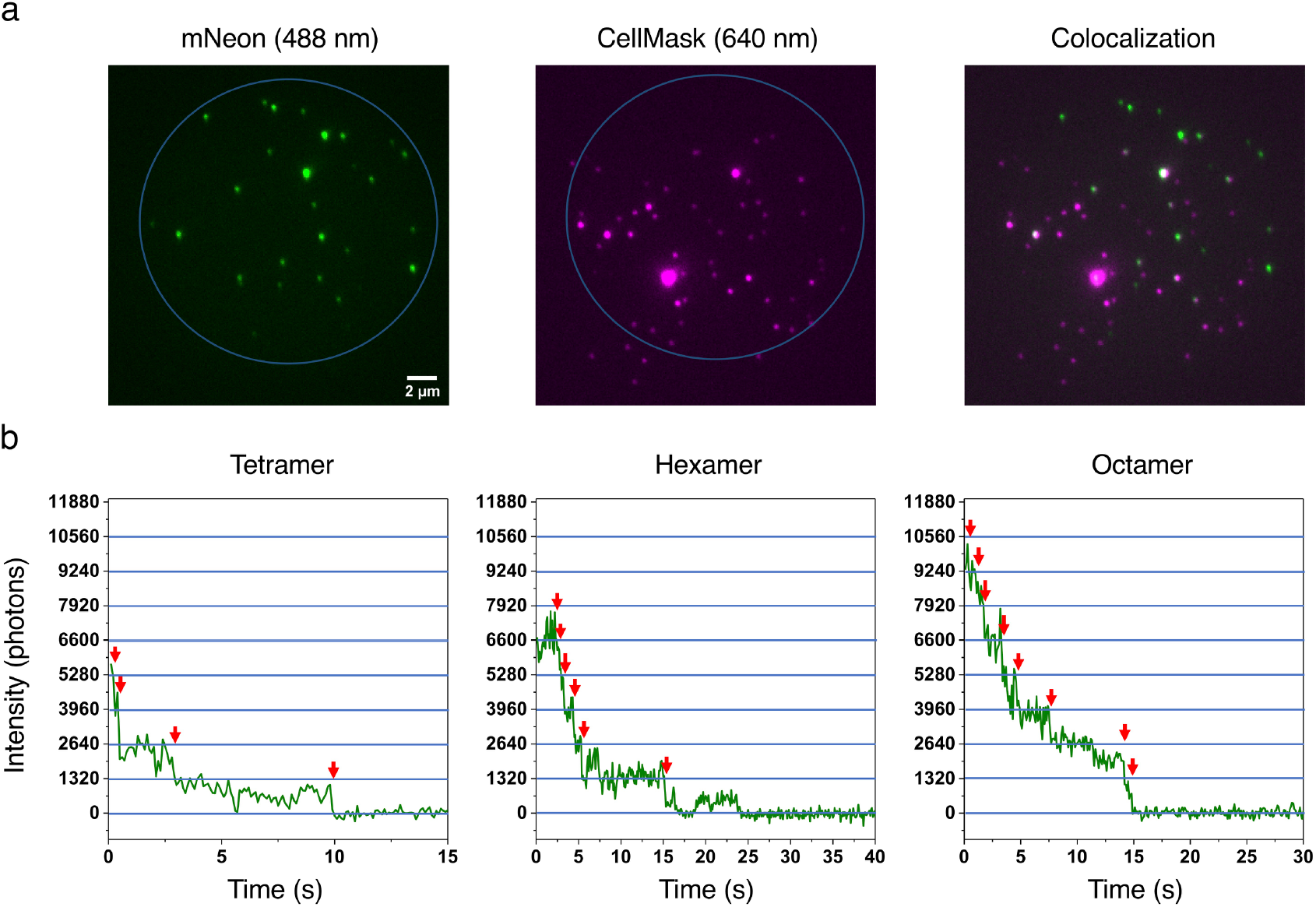
Photobleaching of IMVs with TatABC^mNeon^. **a,** Adherent IMVs. IMVs containing TatABC^mNeon^ stained with the CellMask lipid dye were visualized using 488 nm (mNeon) and 640 nm (CellMask) excitation. The mNeon was imaged first and photobleached before imaging with the red channel. The free CellMask dye does not stick to the coverslip surface (**Supplementary Fig. 4**), indicating that it reliable reports on the presence of membrane lipids. Only co-localized spots (TatC^mNeon^ within an IMV) were used for photobleaching analysis. **b,** Sample photobleaching traces. Steps are established at the times identified by *red* arrows (manually identified). While blinking events occurred and caused deviations from regular steps, the combination of step-size and total intensity allowed assignment of total steps. Individual traces were background subtracted and normalized to zero. **Supplementary Fig. 5** demonstrates the linearity of photon counts with increasing oligomer size and that the average step was 1300 photons (blue lines).

**Fig. 4.**
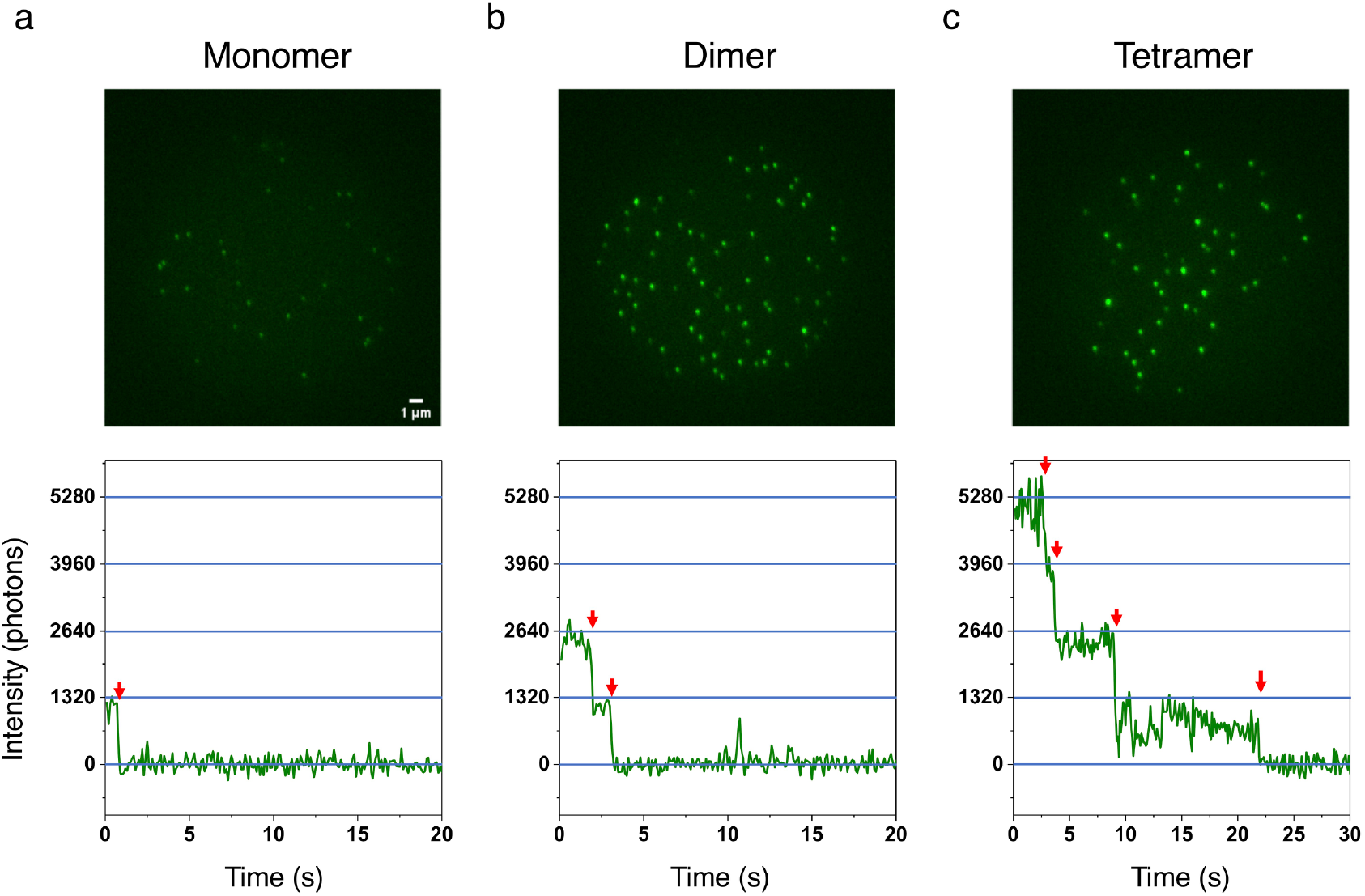
Photobleaching of mNeon Standards. Oligomers of mNeon were imaged and analyzed identically as in **Fig. 3**, except that only 488 nm excitation was used. **a,** Monomer – Cys-free mNeon. **b,** Dimer – the disulfide dimer of mNeonC. **c,** Tetramer – the disulfide dimer of 2xmNeonC.

The step distributions for IMVs containing TatB^mNeon^ or TatC^mNeon^ reveal that all possibilities from 1-8 steps were observed (**Fig. 5**). While > 8 steps were occasionally observed, 8 steps were deemed the limit where manual step-picking was reliable, so anything above this was not included in our analysis. The average step size of mNeon was 1320±20 photons (**Supplementary Fig. 5**). Assuming the simplest model, i.e., that the receptor complex consists of only one TatBC heterodimer, the step histograms were fit assuming that these complexes were Poisson distributed into the IMVs. Since control experiments revealed that ~2.7% of wild-type TatABC IMVs had detectable fluorescence in the mNeon channel, which almost always bleached in a single step (**Supplementary Figure 6**), the 1 step data bin was not included in the fitting routine. While these fits for TatB^mNeon^ or TatC^mNeon^ photobleaching histograms predict an average number of monomeric receptor complexes per IMV of ~3 (**Fig. 5**), it also predicts a low percentage of empty vesicles (~4%), which was clearly not observed (previous paragraph; > 66% of CellMask positive spots were devoid of fluorescent Tat proteins). One possibility is that the Tat receptor complexes were not randomly distributed within the cytoplasmic membrane, which biased how they were distributed into IMVs, thus leading to a higher percentage of empty IMVs than predicted. However, photobleaching histograms for dimeric and tetrameric mNeon exhibited a significant number of molecules exhibiting steps below the nominal oligomer size (**Fig. 6a,b**), which could not be explained by fragments in the protein preparation (**Supplementary Fig. 2**). Thus, a correction involving a photophysical explanation seemed more likely, which is described in the sections that follow.

**Fig. 5.**
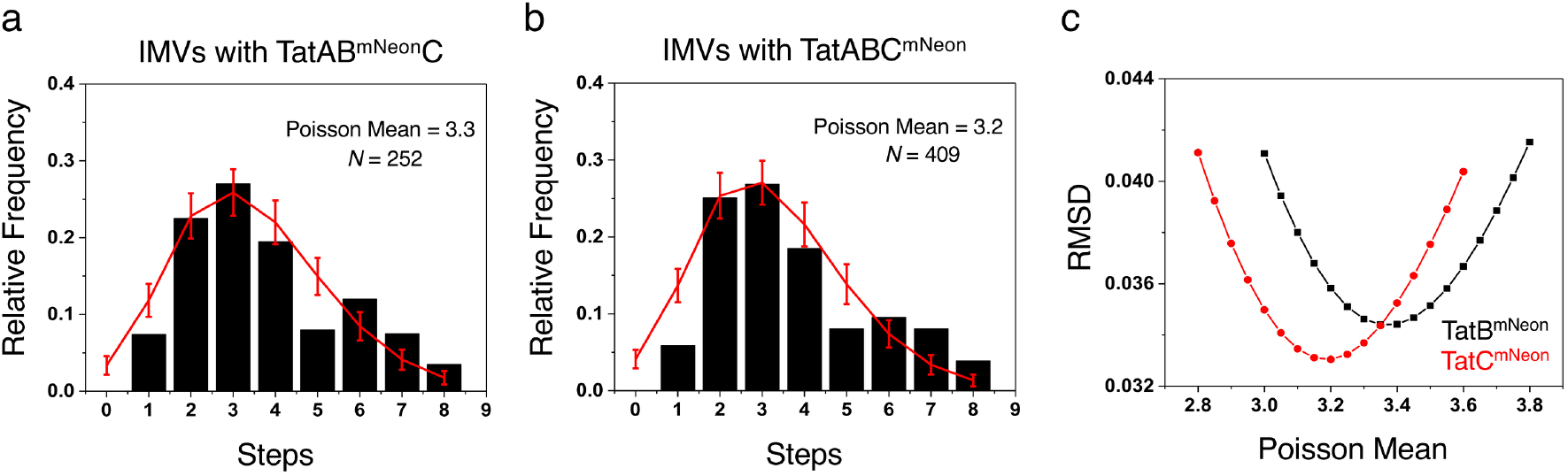
Photobleaching Step Histograms for IMVs with TatAB^mNeon^C or TatABC^mNeon^. **a,b,** Histograms of the number of steps observed in photobleaching profiles of IMVs containing TatAB^mNeon^C or TatABC^mNeon^. The data (*black*) were fit assuming the simplest model, i.e., that receptor complexes contain a single TatBC heterodimer, and that these distributed into IMVs according to a Poisson distribution. The single step bin contained contaminants that bleached in a single step, and occurred at a density of ~2 spots per microscope field (~2.7% of total spots; see text and **Supplementary Figure 6**). Consequently, the data were fit to step bins 2-8. The best fit (*red*) was determined by the least root-mean-square deviation (RMSD) between the experimental and expected values (Poisson probabilities). Error bars are standard deviations (SDs) obtained from 5000 simulated distributions of *N* values each, where *N* is the number of experimental measurements. **c,** The optimal Poisson mean. The RMSD (step bins 2-8) for different Poisson mean values were calculated for the data in **a** and **b**, assuming that all mNeon proteins were actively fluorescent (FDE = 1). The minimum RMSD occurs at Poisson means of ~3.4 and ~3.2 for TatB^mNeon^ and TatC^mNeon^, respectively; the corresponding predicted distributions are shown in **a** and **b**.

**Fig. 6.**
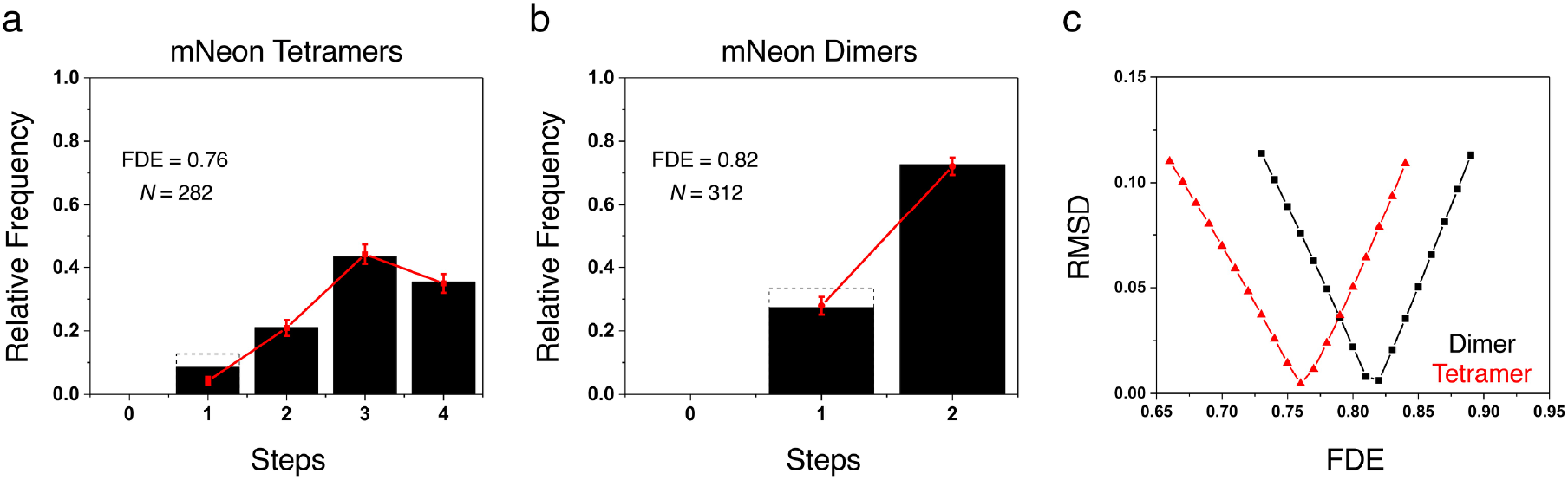
Photobleaching Step Histograms for mNeon Tetramers and Dimers. **a,** Photobleaching step histogram for mNeon tetramers (disulfide dimers of 2xmNeonC). The data (*black*) were fit assuming that the fluorescently active mNeon proteins were binomially distributed within the individual tetramers, yielding a best fit (*red*) fluorescence detection efficiency (FDE) of 0.76. The single step data were ignored for the fit shown, acknowledging the known presence of surface contaminants. The corrected single step data (*solid*; original data shown as *dashed box*) obtained by subtracting the estimated number of contaminants (~1.9 spots/field; see **Supplementary Fig. 4**) agree better with the prediction from the fit. **b,** Photobleaching step histogram for mNeon dimers (disulfide dimers of mNeonC). Assuming a binomial distribution of the fluorescently active mNeon proteins, the corrected data (*solid black*) predict an FDE ≈ 0.82. The correction here (original data shown as *dashed box*) is substantially less than in **a** since the dimer single step data were a larger fraction of the total and the dimer spots were somewhat more densely distributed on the surface than the tetramer spots. **c,** The optimal FDE. The RMSDs for different FDE values were calculated for the data in **a** and **b**: (*red*) tetramer fits to step bins 2-4; (*black*) dimer fits to corrected data.

### Fluorescence Detection Efficiency (FDE) of mNeon

Fluorescent proteins are not always detectable, a phenomenon that is typically assumed to largely arise from deficiencies in fluorophore maturation^49–53^. A non-unity fluorescence detection efficiency (FDE) results in undercounting the number of fluorophores present. For an FDE significantly less than 1, the number of steps corresponding to the nominal oligomer size may not be the most frequent observation (e.g., **Fig. 6a**), thereby complicating the identification of an unknown oligomeric state. Assuming that non-fluorescent monomers were binomially distributed into mNeon tetramers, the FDE of mNeon in this oligomer was estimated as 0.76 (**Fig. 6a,c**). The FDE of mNeon in the dimer was ~0.82 (**Fig. 6b,c**), indicating a higher detection efficiency.

### The Tat Receptor Complex is a Tetramer

The analysis of the IMV photobleaching histograms was more complex than that for the mNeon oligomers since the IMVs contain variable numbers of Tat receptor complexes. We assumed a model in which the number of Tat receptor complexes was Poisson distributed into IMVs, and the number of active fluorophores in each oligomer were binomially distributed, as determined by the FDE. Thus, there were two fitting parameters (Poisson mean and FDE). For both TatB^mNeon^ and TatC^mNeon^, the best fits resulted when the receptor complex was assumed to be a tetramer (**Fig. 7**, **Supplementary Figure 7**, and **Supplementary Table 1**). This conclusion was confirmed with duplicate datasets for each labeled Tat protein (**Supplementary Fig. 8** and **Supplementary Table 1**). Notably, the FDE was significantly lower for the mNeon tag on the Tat proteins (~0.6-0.7, assuming tetramers), indicating both that fusion to the Tat proteins affected fluorophore maturation, and that the FDE must be directly estimated from the experiment rather than from simple control proteins.

**Fig. 7.**
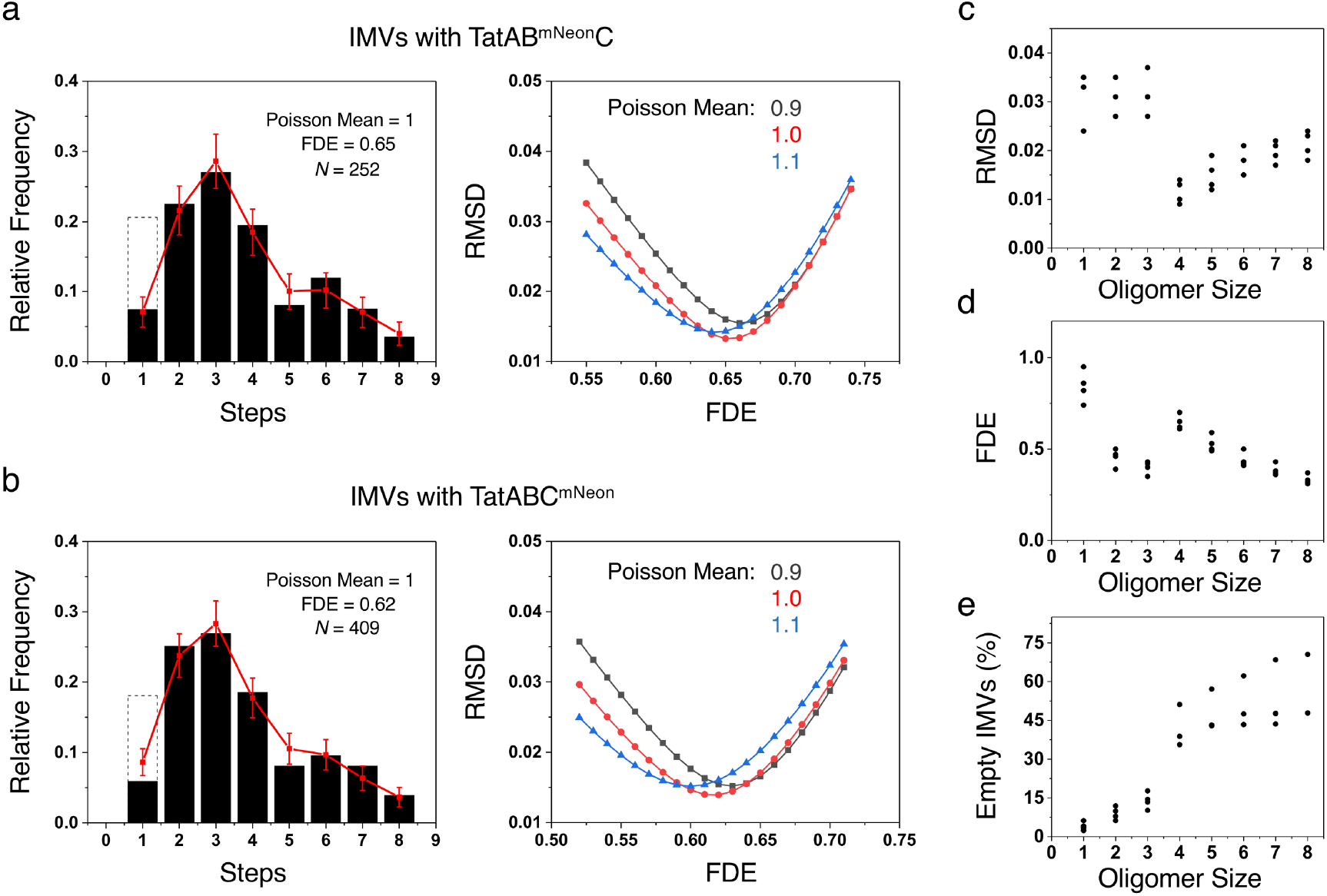
The Tetrameric Tat Receptor Complex. **a,b,** Best fits to photobleaching data for IMVs containing TatAB^mNeon^C or TatABC^mNeon^. By varying the assumed oligomerization state (monomers to octamers), the Poisson mean, and the FDE, the best fits to the experimental photobleaching data for IMVs containing TatAB^mNeon^C (**a**) or TatABC^mNeon^ (**b**) indicate that the Tat receptor complex is a tetramer. Sample fits for other oligomerization states are shown in **Supplementary Fig. 7**. The single step data in the histograms are corrected (*solid black*, see **Fig. 6**); uncorrected values are also indicated (*dashed black boxes*). Only step bins 2-8 were included in the RMSD for the best fits shown (*red*); inclusion of the corrected single step bin for both data sets leads to identical conclusions (see **Supplementary Table 1**). **c,d,e,** Summary of fit parameters and predicted empty IMVs. The RMSDs (**c**), FDEs (**d**), and predicted percentage of empty IMVs (**e**) assuming various oligomerization states for two preparations each of IMVs containing TatAB^mNeon^C or TatABC^mNeon^ (**Fig. 5** and **Supplementary Fig. 8**) show a significant transition between the trimer and tetramer. The best fit (lowest RMSD) was obtained assuming a tetramer (as shown in **a** and **b**). Raw data are summarized in **Supplementary Table 1**.

## DISCUSSION

The reported work probed the oligomerization state of the Tat receptor complex in a native membrane environment by counting photobleaching steps. While conceptually straightforward, this approach typically encounters significant challenges: i) the photobleaching steps must be distinguishable; ii) the interpretation relies on the FDE (which can vary depending on the fusion protein); iii) the number of expected steps is dependent on the number of complexes in an analyzed spot; and iv) potential interference from background components must be accounted for. Point (i) was addressed by using mNeon, a bright, relatively photostable fluorescent protein^43^. While blinking was observed, this did not eliminate our ability to manually count steps. If anything, blinking yielded an underestimated FDE, but not a material change in our major conclusion. Points (ii) and (iii) were addressed by including the FDE and complex distribution (Poisson mean) in the data analysis routine as fitting parameters. The FDE was inversely correlated with polypeptide length for the two mNeon-tagged Tat proteins and the mNeon dimer and tetramer standards, indicating differences in chromophore maturation efficiency for distinct fusion proteins. Background components (point iv) were eliminated by spot selection (e.g., co-localized lipid marker and mNeon fluorescence) or subtraction (e.g., single step photobleaching spots on the coverslips or arising from non-tagged IMVs). Our analysis revealed that a tetrameric Tat receptor complex is the most parsimonious interpretation of the photobleaching data. We first expand on the support for this conclusion and then discuss the implications.

The RMSD between the experimental data and the best fit assuming a given oligomerization model is the primary indication that the Tat receptor complex is a tetramer. Four distinct preparations of IMVs – two containing TatAB^mNeon^C and two containing TatABC^mNeon^ – yielded the same conclusion, namely, that the lowest RMSD was obtained by assuming a tetrameric receptor complex (**Supplementary Table 1**). While there is a clear and significant break between the RMSD for tetramers and lower order oligomers (trimers and lower; **Fig. 7c**), the RMSD for somewhat higher order oligomers were similar to the tetramer fit. However, there are additional considerations.

Maturation efficiencies (including for mNeon) are typically above 0.5^49–51,54–58^, suggesting that dimer, trimer, and hexamer or higher models are unlikely (**Fig. 7d**). A functional analysis of TatC fusion proteins revealed that the TatC oligomerization state must be an even number^59^, thus ruling out the pentamer model (as well as other odd oligomerization states).

The high number of Tat-deficient IMVs also argues against the trimer and lower models. From the colocalization analysis, 67-82% of the lipid containing structures (spots exhibiting CellMask fluorescence) did not contain an mNeon-labeled Tat protein. This is substantially higher than the number of empty vesicles predicted for the trimer and lower oligomerization models, and somewhat to moderately higher than predicted for the tetramer and higher models (**Fig. 7e**). This is not a consequence of low lipid dye concentration, as a 10-fold increase in the concentration of CellMask membrane dye yielded the same colocalization percentage (**Supplementary Fig. 9**). Possible explanations for the lower than expected number of Tat-deficient lipid structures include: i) the Tat receptor complexes were not homogeneously distributed within the bacterial plasma membrane^24,60^; and ii) outer membrane contaminants. Alternatively, some of the receptor complexes could have been released from the membrane by the cell rupture procedure, thus leading to a reduced fraction of IMVs with receptor complexes. The 35-70% of the mNeon Tat fusion proteins not associated with lipid structures is consistent with this interpretation. However, since TatC is a highly hydrophobic membrane protein, only a harsh treatment is expected to remove it from a lipid environment, arguing against this explanation.

The low but measurable transport efficiencies (≤ ~8%) of the IMVs used for photobleaching studies (**Fig. 2a**) suggest that not all of the spots with the colocalized lipid and Tat proteins (≥ 18%) represent transport competent IMVs. In addition to a Tat receptor complex, transport competence requires sufficient TatA and respiratory proteins (to generate the necessary pmf) to partition into the IMV. Since higher TatABC expression levels increase transport competence, TatA is likely the primary limiting protein, although multiple receptor complexes per IMV would also increase transport efficiency. Considering a cell size of ~2 (length) x 1 (width) μm for MC4100 (not shown), ~200 IMVs (100 nm diameter) could be obtained from a single *E. coli* bacterium. A Poisson mean of 1 for the number of complexes per IMV (**Supplementary Table 1**) therefore implies ~200 Tat receptor complexes per *E. coli* cell. Since overproduction of TatABC is necessary to observe in vitro transport with IMVs^7^, this value of 200 is an upper limit for the number of wildtype Tat receptor complexes in non-overproducing cells, likely by a wide margin^20^. In short, the large number of IMVs obtained per *E. coli* cell and the low wildtype levels of Tat proteins indicates that significant Tat protein overproduction levels are required to obtain high overall transport efficiencies as well as transport into most IMVs.

The mechanism of Tat translocation remains under debate. Nonetheless, most recent models place the signal peptide binding site on the inside of the Tat receptor complex^27–29,61,62^. In such a mechanism, cargo translocation would either occur though the center of the receptor complex, or subsequent to a substantial conformational change wherein the necessary conduit is generated in a more external location. In the former case, expansion of the resting pore would be necessary, likely upon recruitment of TatA, as a tetrameric receptor complex is not expected to be able to accommodate all folded protein cargos. In either case, it is unclear if a channel exists in the center of the tetrameric receptor complex, which, if it exists, would necessarily need to be gated closed to prevent collapse of the pmf. Establishing that the Tat receptor complex is a tetramer establishes a critical framework for addressing these structural and functional issues.

## METHODS

### Bacterial strains, growth conditions and plasmids

The *E. coli* strains MC4100ΔTatABCDE (DADE), JM109, and BL21(λDE3) were described earlier^44,63,64^. JM109 was used for clone amplification and plasmid maintenance. BL21(λDE3) cultures producing pre-SufI and the mNeon fluorescent standards from the indicated plasmids were grown in Luria-Bertani (LB) medium^65^ at 37°C. Tat proteins were expressed in the DADE strain and grown in a modified low-salt LB media with glycerol (1% bactotryptone, 0.5% yeast extract, 0.25% NaCl and 0.05% glycerol). All cultures were supplemented with the appropriate antibiotic (30 μg/mL kanamycin or 50 μg/mL ampicillin). All new plasmids were submitted to Addgene, and their construction is described in the history of the linked SnapGene files. Coding sequences were verified by DNA sequencing. Plasmid constructions are briefly outlined below, and the encoded amino acid sequences are indicated in **Supplementary Fig. 1**.

#### pH6-mNeon

The pET-28a_mScarlet plasmid was created by amplifying mScarlet from pCytERM mScarlet N1 (Addgene #85066) with an N-terminal 6xHis-tag and an SpeI site between the 6xHis-tag and mScarlet, and inserting the H6-SpeI-mScarlet PCR product into pET28a (Novagen #69864) using NcoI and BamHI. The mNeon coding sequence amplified from plasmid L40C-CRISPR.EFS.mNeon (Addgene #69146) was then used to replace mScarlet using SpeI and BamHI, generating plasmid pH6-mNeon.

#### pH6-mNeonC

The mNeon coding sequence was amplified from pH6-mNeon using primers that introduced a C-terminal cysteine residue, and the PCR product was then reinserted back into the same plasmid using SpeI and BamHI.

#### pH6-2xmNeon

The mNeon coding sequence was amplified from the pH6-mNeon plasmid with an SpeI site at both ends, and the PCR product was inserted upstream of the existing mNeon gene in the pH6-mNeon plasmid using SpeI, generating plasmid pH6-2xmNeon (correctly oriented product).

#### pH6-2xmNeonC

The plasmid pH6-2xmNeonC was created identically as plasmid pH6-2xmNeon except that the PCR product containing the second mNeon gene was inserted upstream of the existing mNeon copy in the pH6-mNeonC plasmid using SpeI.

#### pTatAB^mNeon^C

The mNeon coding sequence was amplified from the pH6-mNeon plasmid, and the PCR product was used to replace mCherry in pBAD_TatAB^mCherry^C^18^ using KpnI and SpeI, generating plasmid pTatAB^mNeon^C.

#### pTatABC^mNeon^

The pTatABC plasmid^66^ was modified to contain restriction sites after each Tat gene using the QuikChange protocol. The new plasmid (pTatAkBxCs) contained a KpnI restriction site after TatA, an XbaI site after TatB, and a SalI site after TatC. The mNeon coding sequence was amplified from the pH6-mNeon plasmid with a SalI site at both ends, and the PCR product was inserted downstream of TatC in pTatAkBxCs using SalI, generating plasmid pTatABC^mNeon^ (correctly oriented product).

### Protein production and purification

Monomeric and dimeric mNeon were obtained from 500 mL cultures inoculated with 5 mL starter cultures grown for 16 h. Protein production was induced at OD_600_ ≈ 0.6 with 1 mM isopropyl β-D-1-thiogalactopyranoside (IPTG) and growth was continued for 3 hours at 37°C. Cell pellets were stored at −80°C until purification. After frozen cells were thawed on ice, they were rapidly resuspended by adding 40 mL lysis buffer (100 mM CAPS, 250 mM NaCl, 1% Triton X-100, pH 9.5) containing 1X CelLytic B (Cat. #C8740, Sigma), protease inhibitors [10 mM phenylmethylsulfonyl fluoride (PMSF), 20 μg/mL trypsin inhibitor, 10 μg/mL leupeptin and 10 μg/mL pepstatin], 20 μg/mL DNase, and 10 μg/mL RNase. The cell suspension was cleared by centrifugation at 40,000 x *g* for 20 min at 4°C. The supernatant was loaded onto a 10 x 1 cm column with 2 mL Ni-NTA Superflow resin (Cat. #30230, Qiagen) that was pre-equilibrated with Buffer A (10 mM CAPS, 1 M NaCl, 0.1% Triton X-100, pH 9.5). The resin was washed with 30 mL of Buffer A, 30 mL of 10 mM CAPS, 1 M NaCl, pH 9.5, 10 mL of 10 mM CAPS, 0.1 M NaCl, pH 9.5, and 5 mL of 10 mM CAPS, 50 mM NaCl, 50% glycerol, pH 9.5. The bound protein was eluted with 500 mM imidazole, 50 mM NaCl, 50% glycerol, pH 8. Eluates were analyzed by SDS-PAGE to check for purity and concentration. For disulfide dimers of mNeonC and 2xmNeonC (yielding disulfide dimers and tetramers, respectively), the mNeon-containing elution fractions were incubated overnight at 4°C and then purified using size exclusion chromatography with FPLC buffer (50 mM Tris-HCl, 150 mM NaCl, pH 7.5) the next day (Amersham Pharmacia Biotech AKTAdesign system with Superdex 75 column for the disulfide dimer, and Bio-Rad NGC system with Enrich SEC 650 column for the tetramer). The mNeon genetic dimer (2xmNeon) was also purified via size exclusion chromatography using FPLC buffer (Bio-Rad NGC system with Enrich SEC 75 column).

The Tat substrate pre-SufI(IAC) was overproduced from plasmid p-preSufI(IAC) (Addgene #168516). A starter culture (5 mL) grown for 16 h was transferred to a 500 mL culture. Protein production was induced at OD_600_ ≈ 0.6 with 1 mM IPTG, and 25 ml 0.5 M CAPS, pH 10 was added to raise the culture pH^17^. After growth for 2 h at 37°C, cells were recovered (5000 x *g*, 10 min, 4°C) and then stored at −80°C until purification. The pre-SufI(IAC) protein was purified via Ni-NTA chromatography following the same procedure used for the mNeon protein standards. To label with a fluorescent dye, purified pre-SufI(IAC) was first incubated with 1 mM *tris*[2-carboxyethylphosphine] hydrochloride (TCEP) for 10 min, and then with a 50-fold molar excess of Alexa647 maleimide at room temperature (RT) in the dark. After 15 min, the reaction was quenched with 10 mM β-mercaptoethanol (βME). The mixture was diluted 10-fold to reduce the imidazole concentration and then re-purified by Ni-NTA chromatography.

### Isolation of IMVs

IMVs were prepared from the DADE strain transformed with pTatAB^mNeon^C or pTatABC^mNeon^. A starter culture (5 mL) grown overnight at 37°C was used to inoculate 2 L of culture (6×500 mL). Tat protein production was induced at OD_600_ ≈ 0.6 with 0.7% arabinose for 1 h or 4 h at 37°C. Cells were recovered (5000 x *g*, 10 min, 4°C) and then stored at −80°C until the next day. IMVs were isolated as described^18^, except that the three step sucrose gradient used was 2.1 M, 1.5 M and 0.5 M.

### In vitro transport assays

Transport of pre-SufI(IAC)^Alexa647^ into IMVs was assayed essentially as described previously^7,18,46^. In short, transport reactions (35 μL) included IMVs (OD_600_ = 5.0) and pre-SufI(IAC)^Alexa647^ (50 nM) in Transport Buffer (TB; 25 mM MOPS, 25 mM MES, 5 mM MgCl_2_, 50 mM KCl, 200 mM sucrose, pH 8.0) with 1 μg/μL of BSA. Transport of substrate into the IMVs was initiated by the addition of 4 mM NADH, and continued for 20 min at 37°C. Untransported (external) substrate was digested by incubating with 0.57 μg/μL proteinase K for 30 min at RT. Digestions were quenched by the addition of PMSF (5 mM) and SDS Sample Buffer (100 mM Tris-HCl, 4.5 M Urea, 2% SDS, 5% glycerol, 0.02% bromophenol blue, 0.2% βME, pH 6.8). Samples were heated at 95°C for 10 min and then analyzed on an 12% SDS-PAGE gel.

### Western blotting

Western blots were performed as described earlier^67^. TatB and TatC were detected using anti-TatB and anti-TatC primary antibodies^66^ (1:2500 dilution), and polyclonal goat anti-rabbit IgG-HRP secondary antibodies (1:10,000 dilution; Santa Cruz Biotech, Inc).

### Protein analytical methods

Concentrations of purified proteins were determined by the densitometry of bands on Coomassie Blue R-250 stained SDS-PAGE gels using carbonic anhydrase as a standard and a ChemiDoc MP imaging system (Bio-Rad Laboratories). Fluorescent proteins were detected by direct in-gel fluorescence imaging using the same ChemiDoc imaging system. Alexa647 and mNeon concentrations were determined using ε_650_ = 270,000 cm^−1^ M^−1^ and ε_506_ = 116,000 cm^−1^ M^−1^,^43^ respectively. Western blot bands were visualized by chemiluminescence using the Clarity Max Western blotting kit (Bio-Rad Laboratories, #1705062) and the ChemiDoc imaging system. IMV concentrations were determined as the *A*_280_ in 2% SDS^7^.

### Dynamic light scattering (DLS)

DLS experiments were performed on Zetasizer Nano (Malvern Panalytical). IMV samples were diluted 1:100 with 50 mM Tris, 150 mM NaCl, pH 7.5 for analysis.

### Coverslip Preparation

Coverslips (24 mm x 60 mm; VWR, #16004-312) were cleaned by sonicating in distilled water for 10 min, followed by sonicating in acetone for 10 min. After air-drying in a hood, the coverslips were plasma cleaned in argon for 5 min on the high setting (Harrick Plasma, Model PDC-001). The plasma cleaned slides were mounted in home-machined aluminum microscope holders and held in place by high-vacuum grease (Dow Corning, VWR #59344-055). Flow chambers were constructed from strips of double-sided tape and a top coverslip (10.5 mm x 22 mm; Electron Microscopy Sciences, #72191-22).

### Microscopy

Single particle photobleaching analyses were performed with a Nikon Eclipse Ti using a 100X oil-immersion objective (Nikon Apo TIRF, 1.49 NA) and narrowfield epifluorescence excitation^48^ with 488 and 640 nm diode lasers (Coherent). Images were magnified (1.5X tube lens) and recorded using an EMCCD (Evolve Delta; Photometrics) or CMOS (Prime 95B; Photometrics) camera. The focus was adjusted, focus-lock engaged, and the coverslip translated such that the recordings for the samples analyzed captured the photobleaching profile from the moment samples were first illuminated. This insured that no photobleaching events were missed. Exposures were 100 ms with a recording length of 500-600 frames.

All buffers that were used in the single particle imaging protocol were passed through a 0.22 μm syringe filter, which substantially reduced the number of fluorescent background particles. The purified mNeon standards were diluted with Dilution Buffer (50 mM Tris-HCl, 150 mM NaCl, pH 7.5), incubated in flow chambers for a minimum of 5 min, washed with two lane volumes of Dilution Buffer (~20 μL), and then imaged. IMVs (OD_600_ = 5.0) containing TatB^mNeon^ and TatC^mNeon^ were first incubated with 0.1X CellMask Deep Red Plasma Membrane Stain (ThermoFisher Scientific, # C10046) for 15 min at RT. The labeled IMVs were then diluted with Dilution Buffer, incubated in a flow chamber for a minimum of 5 min, washed with two lane volumes of Dilution Buffer, and then imaged.

### Image analysis

Fluorescent spots in photobleaching videos were selected manually using ROI Manager in the Fiji package of ImageJ^68^. For the mNeon standards, all spots were selected. For IMVs, only co-localized spots were selected (spots visualized in both the 488 and 640 nm excitation channels). For every 10×10 pixel ROI chosen, an adjacent area of the same size with no fluorescence was chosen for background subtraction. Intensities were obtained via the multi-measure function of ROI Manager.

Photobleaching profiles were plotted using MATLAB, and steps were identified manually.

### Simulations

Step expectation values were calculated assuming that Tat receptor complexes partitioned into IMVs following a Poisson distribution and the active fluorophores within receptor complexes and mNeon dimer and tetramer standards followed a binomial distribution, where the FDE defines the probability that the fluorophore is actively fluorescent (**Supplementary Software Files 1** and **2**). Standard deviations were obtained from 5000 independent trials of *N* measurements (where *N* = number of experimentally spots analyzed) using Microsoft Excel with the RiskAMP Monte Carlo plugin (Structured Data LLC) (**Supplementary Software File 3**).

## ACKNOWLEDGEMENTS

We thank T.L. Yahr for antibodies to TatB and TatC, T. Palmer for the DADE strain, S. Hamsanathan for assistance with preliminary studies, and B. Pettijohn for making the parent pTatA_k_B_x_C_s_ plasmid. This research was supported by the National Institutes of Health (GM116995 to SMM).

## DATA AND CODE AVAILABILITY STATEMENT

Plasmids pET-28a_mScarlet, pH6-mNeon, pH6-mNeonC, pH6-2xmNeon, pH6-2xmNeonC, pBAD_TatAB^mCherry^C, pTatAB^mNeon^C, pTatA_k_B_x_C_s_, and pTatABC^mNeon^ are available from Addgene (IDs# 178460-178465, 178543-45). The authors declare that all other data supporting the findings of this study are available within the paper and its supplementary information files. Raw data and materials are available from the corresponding author upon reasonable request.

## AUTHOR CONTRIBUTION STATEMENT

SMM conceived the approach, AS purified materials, collected and analyzed data, and wrote the manuscript, RC helped with microscope experiments and data analysis, SMM developed the simulations, provided advice, analyzed data, and edited the manuscript.

