## Supplementary Information for "Oligomerization state of the functional bacterial twin arginine translocation (Tat) receptor complex"

**for**

**SUPPLEMENTARY TABLE 1. Summary of best fit values obtained for photobleaching histograms assuming different oligomerization states of the Tat receptor complex**[1]

**TatB$^{mNeon}$:**

| | Poisson Mean | FDE | RMSD | | Predicted Empty IMVs (%) |
|---|---|---|---|---|---|
| | | | Steps 2-8 | Steps 1-8 | |
| **Monomers** | 3.9 | 0.86 | 0.035 | 0.034 | 3.5 |
| | 3.9 | 0.95 | 0.035 | 0.038 | 2.4 |
| **Dimers** | 3.6 | 0.46 | 0.031 | 0.040 | 7.9 |
| | 3.9 | 0.47 | 0.035 | 0.048 | 6.2 |
| **Trimers** | 2.5 | 0.43 | 0.031 | 0.044 | 13.4 |
| | 2.9 | 0.42 | 0.037 | 0.053 | 10.2 |
| **Tetramers** | 1.0 | 0.65 | 0.013 | 0.012 | 38.8 |
| | 1.1 | 0.70 | 0.010 | 0.010 | 35.6 |
| **Pentamers** | 0.9 | 0.53 | 0.019 | 0.017 | 42.9 |
| | 0.9 | 0.59 | 0.013 | 0.015 | 43.2 |
| **Hexamers** | 0.9 | 0.43 | 0.021 | 0.023 | 43.3 |
| | 0.8 | 0.50 | 0.018 | 0.021 | 47.5 |
| **Heptamers** | 0.9 | 0.38 | 0.022 | 0.024 | 43.6 |
| | 0.8 | 0.43 | 0.021 | 0.025 | 47.7 |
| **Octamers** | 0.8 | 0.33 | 0.024 | 0.026 | 47.8 |
| | 0.8 | 0.37 | 0.023 | 0.029 | 47.8 |

**TatC$^{mNeon}$:**

| | Poisson Mean | FDE | RMSD | | Predicted Empty IMVs (%) |
|---|---|---|---|---|---|
| | | | Steps 2-8 | Steps 1-8 | |
| **Monomers** | 3.9 | 0.82 | 0.033 | 0.039 | 4.1 |
| | 3.8 | 0.74 | 0.024 | 0.060 | 6.2 |
| **Dimers** | 3.1 | 0.50 | 0.027 | 0.049 | 9.9 |
| | 3.4 | 0.39 | 0.027 | 0.074 | 11.9 |
| **Trimers** | 2.5 | 0.40 | 0.027 | 0.052 | 14.4 |
| | 2.4 | 0.35 | 0.031 | 0.080 | 17.7 |
| **Tetramers** | 1.0 | 0.62 | 0.014 | 0.016 | 38.8 |
| | 0.7 | 0.61 | 0.009 | 0.030 | 51.1 |
| **Pentamers** | 0.9 | 0.50 | 0.016 | 0.022 | 43.0 |
| | 0.59 | 0.49 | 0.012 | 0.037 | 57.1 |
| **Hexamers** | 0.9 | 0.41 | 0.018 | 0.029 | 43.4 |
| | 0.5 | 0.42 | 0.015 | 0.040 | 62.2 |
| **Heptamers** | 0.8 | 0.36 | 0.019 | 0.030 | 47.6 |
| | 0.4 | 0.37 | 0.017 | 0.042 | 68.4 |
| **Octamers** | 0.8 | 0.31 | 0.020 | 0.034 | 47.9 |
| | 0.37 | 0.32 | 0.018 | 0.046 | 70.5 |

[1]The two rows for each oligomer size are the values obtained for two different preparations of IMVs containing TatAB$^{mNeon}$C or TatABC$^{mNeon}$.

***H6-mNeon***
***(plasmid pH6-mNeon; Addgene #178460):***
MGHHHHHHTSMVSKGEEDNMASLPATHELHIFGSINGVDFDMVGQGTGNPNDGYEELNLKSTKGDLQFSPWILVPHI
GYGFHQYLPYPDGMSPFQAAMVDGSGYQVHRTMQFEDGASLTVNYRYTYEGSHIKGEAQVKGTGFPADGPVMTNSLT
AADWCRSKKTYPNDKTIISTFKWSYTTGNGKRYRSTARTTYTFAKPMAANYLKNQPMYVFRKTELKHSKTELNFKEW
QKAFTDVMGMDELYK

***H6-mNeon-C***
***(plasmid pH6-mNeonC; Addgene #178461):***
MGHHHHHHTSMVSKGEEDNMASLPATHELHIFGSINGVDFDMVGQGTGNPNDGYEELNLKSTKGDLQFSPWILVPHI
GYGFHQYLPYPDGMSPFQAAMVDGSGYQVHRTMQFEDGASLTVNYRYTYEGSHIKGEAQVKGTGFPADGPVMTNSLT
AADWCRSKKTYPNDKTIISTFKWSYTTGNGKRYRSTARTTYTFAKPMAANYLKNQPMYVFRKTELKHSKTELNFKEW
QKAFTDVMGMDELYKC

***H6-2xmNeon***
***(plasmid pH6-2xmNeon; Addgene #178462):***
MGHHHHHHTSGTMVSKGEEDNMASLPATHELHIFGSINGVDFDMVGQGTGNPNDGYEELNLKSTKGDLQFSPWILVP
HIGYGFHQYLPYPDGMSPFQAAMVDGSGYQVHRTMQFEDGASLTVNYRYTYEGSHIKGEAQVKGTGFPADGPVMTNS
LTAADWCRSKKTYPNDKTIISTFKWSYTTGNGKRYRSTARTTYTFAKPMAANYLKNQPMYVFRKTELKHSKTELNFK
EWQKAFTDVMGMDELYKQLGSGSTSMVSKGEEDNMASLPATHELHIFGSINGVDFDMVGQGTGNPNDGYEELNLKST
KGDLQFSPWILVPHIGYGFHQYLPYPDGMSPFQAAMVDGSGYQVHRTMQFEDGASLTVNYRYTYEGSHIKGEAQVKG
TGFPADGPVMTNSLTAADWCRSKKTYPNDKTIISTFKWSYTTGNGKRYRSTARTTYTFAKPMAANYLKNQPMYVFRK
TELKHSKTELNFKEWQKAFTDVMGMDELYK

***H6-2xmNeon-C***
***(plasmid pH6-2xmNeonC); Addgene #178463):***
MGHHHHHHTSGTMVSKGEEDNMASLPATHELHIFGSINGVDFDMVGQGTGNPNDGYEELNLKSTKGDLQFSPWILVP
HIGYGFHQYLPYPDGMSPFQAAMVDGSGYQVHRTMQFEDGASLTVNYRYTYEGSHIKGEAQVKGTGFPADGPVMTNS
LTAADWCRSKKTYPNDKTIISTFKWSYTTGNGKRYRSTARTTYTFAKPMAANYLKNQPMYVFRKTELKHSKTELNFK
EWQKAFTDVMGMDELYKQLGSGSTSMVSKGEEDNMASLPATHELHIFGSINGVDFDMVGQGTGNPNDGYEELNLKST
KGDLQFSPWILVPHIGYGFHQYLPYPDGMSPFQAAMVDGSGYQVHRTMQFEDGASLTVNYRYTYEGSHIKGEAQVKG
TGFPADGPVMTNSLTAADWCRSKKTYPNDKTIISTFKWSYTTGNGKRYRSTARTTYTFAKPMAANYLKNQPMYVFRK
TELKHSKTELNFKEWQKAFTDVMGMDELYKC

***TatB-mNeon***
***(encoded in plasmid pTatAB$^{mNeon}$C; Addgene #178464):***
VFDIGFSELLLVFIIGLVVLGPQRLPVAVKTVAGWIRALRSLATTVQNELTQELKLQEFQDSLKKVEKASLTNLTPE
LKASMDELRQAAESMKRSYVANDPEKASDEAHTIHNPVVKDNEAAHEGVTPAAAQTQASSPEQKPETTPEPVVKPAA
DAEPKTAAPSPSSSDKPYTRVPMVSKGEEDNMASLPATHELHIFGSINGVDFDMVGQGTGNPNDGYEELNLKSTKGD
LQFSPWILVPHIGYGFHQYLPYPDGMSPFQAAMVDGSGYQVHRTMQFEDGASLTVNYRYTYEGSHIKGEAQVKGTGF
PADGPVMTNSLTAADWCRSKKTYPNDKTIISTFKWSYTTGNGKRYRSTARTTYTFAKPMAANYLKNQPMYVFRKTEL
KHSKTELNFKEWQKAFTDVMGMDELYK

***TatC-mNeon***
***(encoded in plasmid pTatABC$^{mNeon}$; Addgene #178465):***
MSVEDTQPLITHLIELRKRLLNCIIAVIVIFLCLVYFANDIYHLVSAPLIKQLPQGSTMIATDVASPFFTPIKLTFM
VSLILSAPVILYQVWAFIAPALYKHERRLVVPLLVSSSLLFYIGMAFAYFVVFPLAFGFLANTAPEGVQVSTDIASY
LSFVMALFMAFGVSFEVPVAIVLLCWMGITSPEDLRKKRPYVLVGAFVVGMLLTPPDVFSQTLLAIPMYCLFEIGVF
FSRFYVGKGRNREEENDAEAESEKTEEVDMVSKGEEDNMASLPATHELHIFGSINGVDFDMVGQGTGNPNDGYEELN
LKSTKGDLQFSPWILVPHIGYGFHQYLPYPDGMSPFQAAMVDGSGYQVHRTMQFEDGASLTVNYRYTYEGSHIKGEA
QVKGTGFPADGPVMTNSLTAADWCRSKKTYPNDKTIISTFKWSYTTGNGKRYRSTARTTYTFAKPMAANYLKNQPMY
VFRKTELKHSKTELNFKEWQKAFTDVMGMDELYK

**Supplementary Figure 1. Protein sequences for the mNeon proteins used in this study.** The mNeon sequences are indicated in *green*, additions/linkers are in *red*, 6xHis tags are in *blue* and the TatB and TatC sequences are *underlined*.

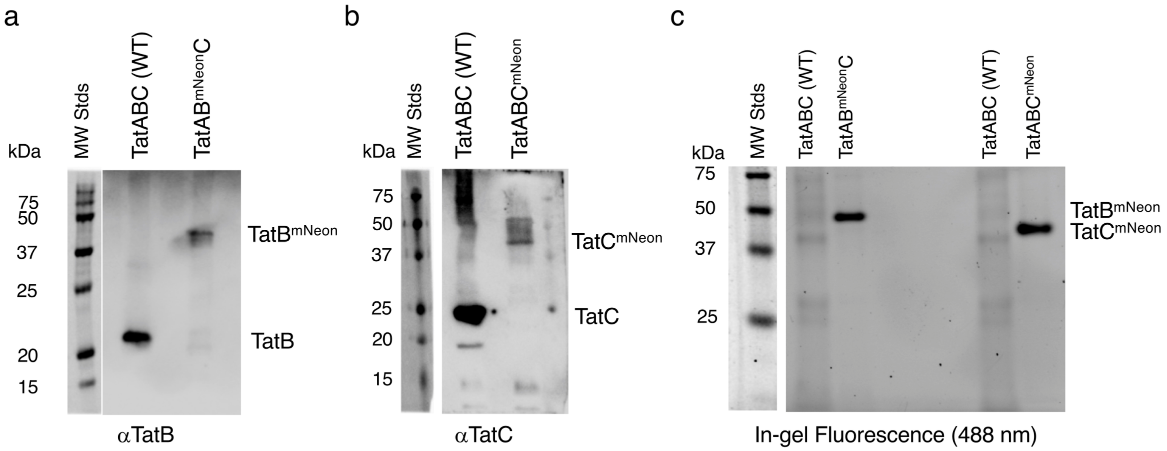


**Supplementary Figure 2. Stability of TatB$^{mNeon}$ and TatC$^{mNeon}$ Fusions. a,b,** Western blots of IMVs containing purified TatAB$^{mNeon}$C and TatABC$^{mNeon}$ were probed with anti-TatB (1:4000) (**a**) and anti-TatC (1:1250) (**b**) antibodies. The shift of ~25 kDa in the TatB$^{mNeon}$- and TatC$^{mNeon}$-containing lanes relative to the bands in the wild-type TatABC lanes corresponds to the molecular weight of mNeon. No free TatB or TatC was observed for the mNeon-tagged proteins, indicating that the mNeon fusion proteins were not subject to proteolysis. **b,** In-gel fluorescence of IMVs containing purified TatAB$^{mNeon}$C and TatABC$^{mNeon}$. Fluorescence bands ($\lambda$ = 488 nm) were detected at a molecular weight consistent with TatB$^{mNeon}$ and TatC$^{mNeon}$ (compare with **a**). The absence of free mNeon confirms that the mNeon fusion proteins were not subject to proteolysis.

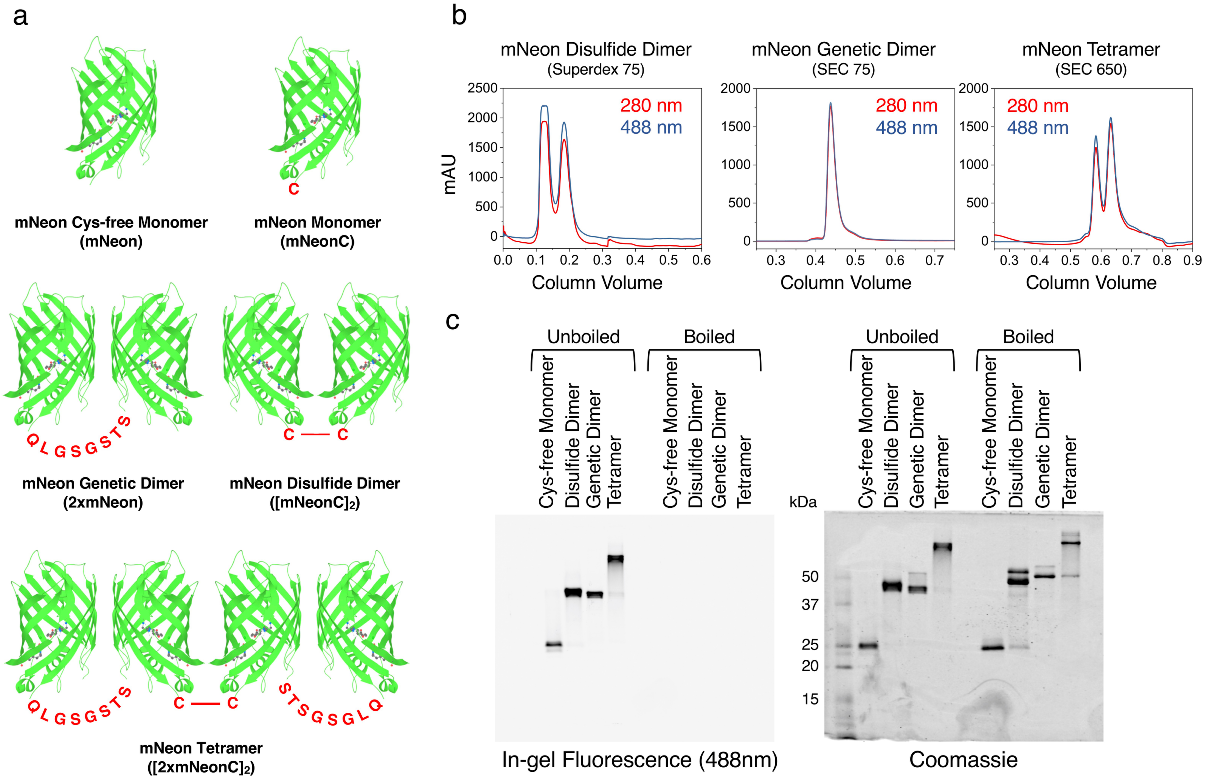


**Supplementary Figure 3. Purification of mNeon Standards. a,** The mNeon standards. The N-terminal 6xHis-tags are not shown (see **Supplementary Fig. 1**). The disulfide dimer was generated by a disulfide bond between mNeon monomers with a C-terminal cysteine. The tetramer was generated by a disulfide bond between two genetic dimers with a C-terminal cysteine. **b,** Gel filtration chromatograms. The columns used are indicated. **c,** In-gel fluorescence and Coomassie stained SDS-PAGE gel images of purified mNeon standards. The monomer was purified by Ni-NTA alone; dimers and the tetramer were further purified by gel exclusion chromatography (**b**). Boiling destroys the mNeon fluorescence. The difference of ~25 kDa between monomer and dimer corresponds to the molecular weight of mNeon.

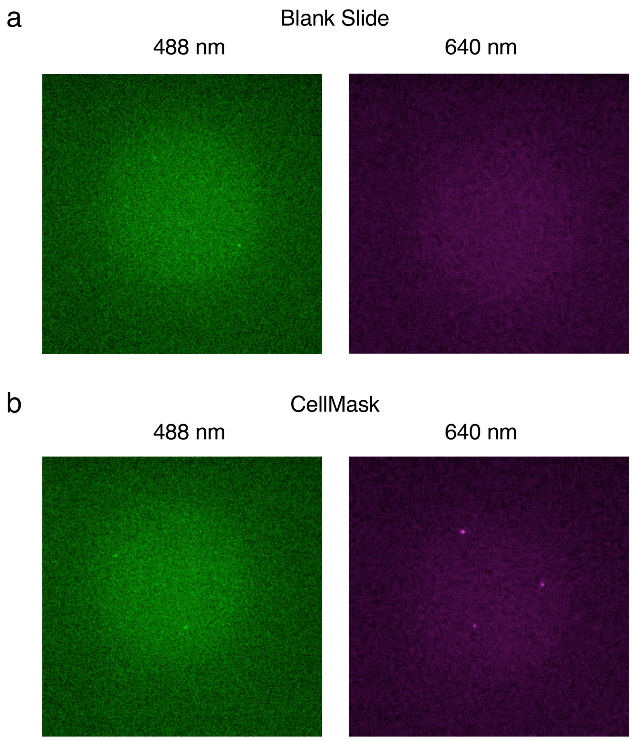


**Supplementary Figure 4. Fluorescence Background. a,** Fluorescence observed for a plasma-cleaned coverslip covered with Dilution Buffer. An average of 1.9 spots/field were observed with $\lambda_{ex}$ = 488 nm that photobleached in a single step. Zero spots/field were observed at $\lambda_{ex}$ = 640 nm. **b**, Fluorescence observed for a plasma-cleaned coverslip treated with 0.1x CellMask dye. The CellMask dye was added in Dilution Buffer, incubated for 15 min at RT, and then washed away. An average of 1.9 and 2 spots/field were observed at $\lambda_{ex}$ = 488 nm and $\lambda_{ex}$ = 640 nm, respectively.

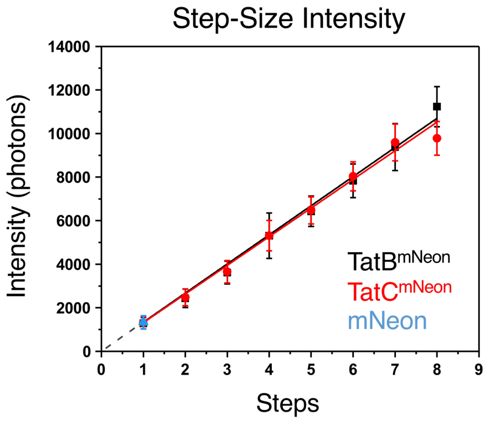


**Supplementary Figure 5. Linearity of Mean Photobleaching Step-size Intensities for IMVs Containing TatAB$^{mNeon}$C and TatABC$^{mNeon}$.** For comparison, the mean intensity obtained for Cys-free mNeon monomers is shown. The error bars (SDs) reveal a significant variation in total intensities, though the steps themselves were identifiable (**Figs. 3** and **4**). Data were fit with a line passing through 0, yielding 1340±20 and 1300±20 photons/step (mean ± standard error) for TatB$^{mNeon}$ and TatC$^{mNeon}$, respectively.

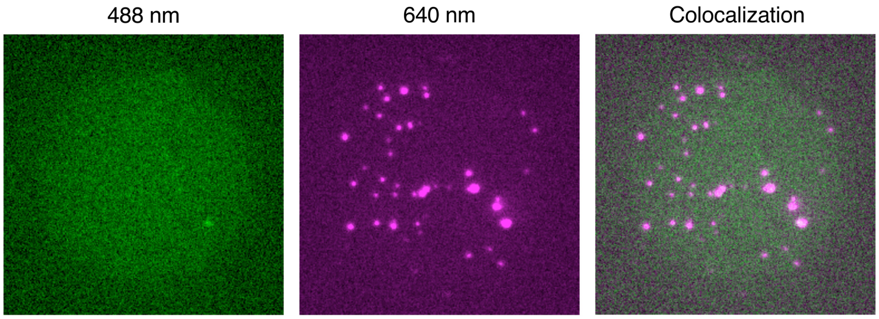


**Supplementary Figure 6. Imaging of IMVs Containing Wildtype TatABC Labeled with the CellMask Dye.** Approximately 2.7% of IMVs were visible with 488 nm excitation. As these spots almost invariably photobleached in a single step, they contaminated the single step histograms in the photobleaching analysis. The data in **Figs. 6,7** and **Supplementary Figs. 7,8** were corrected based on this background contamination.

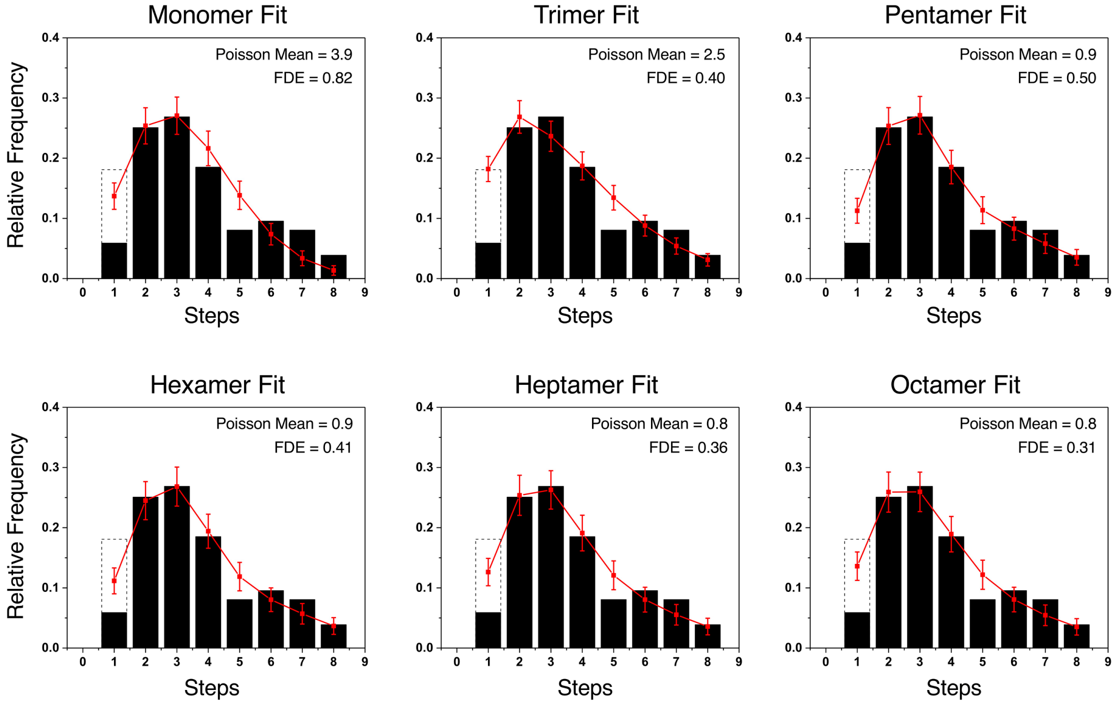


**Supplementary Figure 7. Alternate Fits Assuming Different Oligomeric Sizes for the Tat Receptor Complex.** While a tetramer model for the Tat receptor complex yielded the best fit to the data in **Fig. 5b**), the fits for alternate oligomeric models are shown here (*N* = 409). Fit parameters are summarized in **Figs. 7c,d,e** and **Supplementary Table 1**.

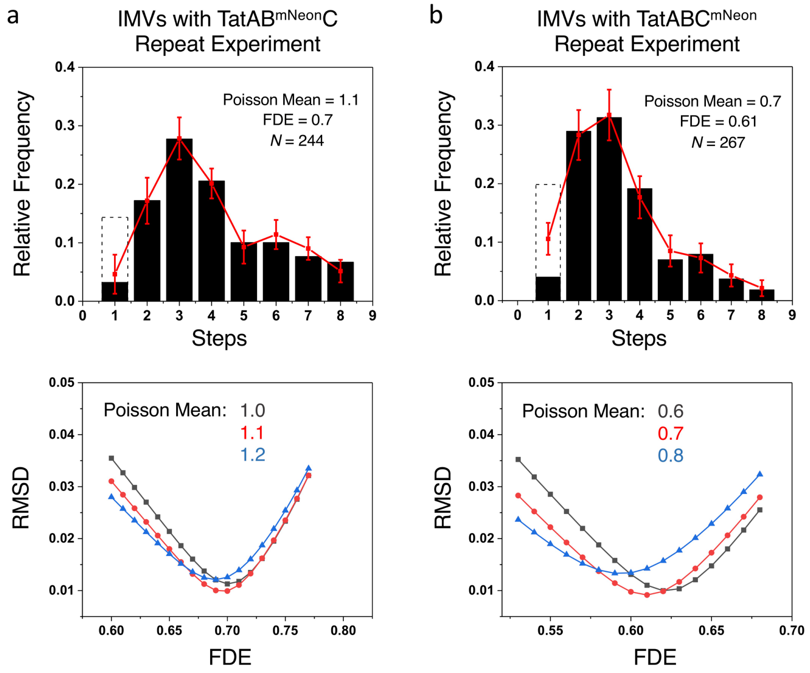


**Supplementary Figure 8. Repeat Photobleaching Experiments of IMVs Containing TatAB$^{mNeon}$C or TatABC$^{mNeon}$.** Independent (duplicate) datasets for the experiments reported in **Fig. 7a,b** (different IMV preparations). Fits for a tetramer model are shown here. Parameters for the different oligomeric fits are reported in **Fig. 7c,d,e** and **Supplementary Table 1**.

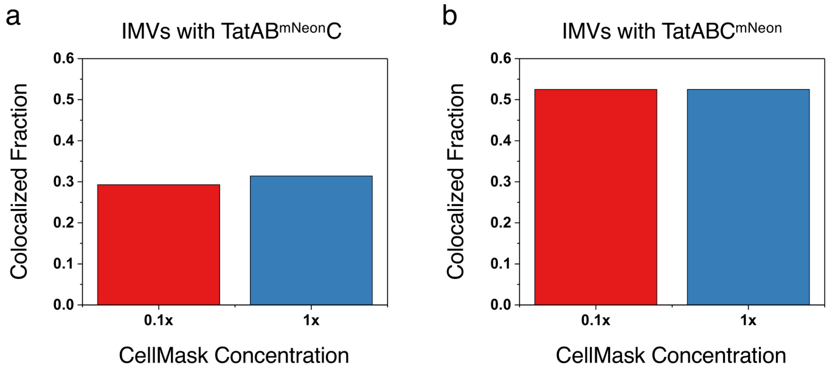


**Supplementary Figure 9. IMV Membrane Labeling Efficiency with CellMask. a,b,** IMVs containing TatAB$^{mNeon}$C (**a**) and TatABC$^{mNeon}$ (**b**) were incubated with two different concentrations of CellMask (15 min at RT). The number of mNeon spots that colocalized with the membrane dye did not significantly change upon increasing the concentration of CellMask membrane dye by 10-fold. The lower (0.1x) concentration was used for all photobleaching experiments.
